# Resistance training-induced appendicular lean tissue mass changes are largely unrelated to pre-training bone characteristics in a larger cohort of untrained adults

**DOI:** 10.1101/2025.05.27.656407

**Authors:** Dakota R. Tiede, Daniel L. Plotkin, Mason C. McIntosh, J. Max Michel, Kevin W. Huggins, Darren T. Beck, Michael D. Goodlett, Joshua C. Carr, Brad J. Schoenfeld, Christopher B. Mobley, Kaelin C. Young, Paul A. Swinton, Andrew D. Frugé, Michael D. Roberts

**Author notes:** Address correspondence to: Michael D. Roberts, PhD, Endowed Alumni Professor, Director, Nutrabolt Applied and Molecular Physiology Laboratory, School of Kinesiology, Auburn University, 301 Wire Road, Office 286, Auburn, AL 36849.

## Abstract

We sought to determine if pre-intervention bone characteristics measured by dual-energy x-ray absorptiometry (DXA) were associated with changes in bone-free lean tissue mass following a period of resistance training in a large cohort of untrained adults (n=119, 62M/57F, 26.0±4.7 kg/m^2^, age range = 18-70 years old). Participants completed 10-12 weeks of supervised whole-body resistance training twice weekly, and DXA scans were obtained approximately the same time of day prior to the intervention and 48-72 hours following the final training bout. Associations between baseline skeletal measures (e.g., appendicular bone characteristics, shoulder and hip widths) and training induced changes in appendicular lean mass were examined by estimating correlations between participant-level random slopes (reflecting change over time) and baseline skeletal measures. The same approach was used to evaluate associations between other participant attributes (e.g., age, training volume-load, self-reported energy intake) and appendicular lean tissue mass changes. Modeling was also used to explore whether baseline skeletal characteristics (e.g., shoulder and hip widths) moderated the change in appendicular lean tissue mass from training. All analyses used a Bayesian framework, and interpretation focused on estimated effect sizes and their associated credible intervals rather than formal null hypothesis testing. Strong positive associations were observed between pre-intervention characteristics including dual-arm lean tissue mass and dual-arm bone mineral content (*r*=0.90), dual-leg lean tissue mass and dual-leg bone mineral content (*r*=0.86), dual-leg lean tissue mass and pelvic mineral content (*r*=0.73), and dual-arm lean tissue mass and shoulder width (*r*=0.76). In contrast, weak associations were observed between training-induced changes in appendicular lean tissue mass versus bone characteristics, training volume-load, self-reported energy intake, self-reported protein intake, BMI, and age (−0.08≤*r*≤ 0.24). After adjusting for sex, multivariable analyses indicated minimal evidence that skeletal characteristics moderated the hypertrophic response to training. These findings do not support a meaningful role of pre-training bone characteristics in influencing the lean tissue mass adaptations to shorter-term resistance training.

## INTRODUCTION

Research indicates correlations between certain bone measures (e.g., size and geometry) and skeletal muscle mass across different populations (1-4). This relationship is bidirectional and has several mechanistic bases. For instance, bone and muscle tissue share a common mesenchymal stem cell lineage during embryogenesis (5). Both tissues are multinucleated and experience rapid growth phases during adolescence through the growth hormone/insulin-like growth factor-1 axis (6). Endocrine signaling between tissues facilitates crosstalk via osteokines and myokines (7). This interdependence is particularly apparent with advancing age where the presence of sarcopenia is associated with an increased risk of osteopenia (8), underscoring the shared adaptability of these tissues to mechanical loading/unloading. In fact, three theoretical frameworks have been used to describe the muscle-bone mechanical coupling phenomena including Wolff’s law (i.e., bone adapts through mechanical stress), the Utah paradigm (i.e., bone adapts through mechanical, cellular, and biochemical factors), and the mechanostat hypothesis (i.e., bone adapts through biological feedback dictated by mechanical strain) (9). Skeletal muscle is responsive to a similar host of external stimuli, although through distinct mechanisms and an altered time-course (10, 11).

Building on this bone-muscle relationship, several studies have reported that months-to-years of resistance training concomitantly increase bone mineral density and skeletal muscle mass (12-14). While it is plausible to assume that bone characteristics (i.e., size and density) may influence the hypertrophic response to resistance training, this relationship has not been well explored. This potential link between tissues seems practical given that a strong skeletal system is likely better equipped to physically support skeletal muscle hypertrophy. A report by van Etten et al. (15) provides some support for this association. These researchers examined the hypertrophic response heterogeneity to 12 weeks of resistance training in younger adult males and reported that those having a “slender” body build did not experience increases in fat-free mass whereas those considered “solid” gained 1.6 kg on average. Though the authors contended that “…*muscle, fat, and bone are the three major structural components that model the human body build*”, body composition was assessed using hydrostatic weighing, thus precluding the determination as to how bone characteristics may have influenced lean tissue mass changes between cohorts.

Given this knowledge gap, the primary aim of this study was to determine if pre-intervention bone characteristics (determined using dual-energy x-ray absorptiometry [DXA]) were associated with changes in bone-free lean tissue mass following 10-12 weeks of resistance training in pooled cohorts of younger and older adults. For this primary aim, we anticipated that pre-intervention appendicular bone characteristics (i.e., bone areas, bone masses, and the widths of hips and shoulders defined by skeletal landmarks) would be strongly positively associated with changes in appendicular bone-free lean tissue mass following resistance training. Secondary analyses were also performed to determine if baseline appendicular lean tissue mass, volume-load, self-reported energy and protein intakes, and/or age were associated with resistance training responses. Lastly, multivariable models were constructed to assess whether combinations of skeletal characteristics, baseline muscularity, training variables, dietary intake, and demographic factors jointly explained variance in lean tissue adaptations. We did not adopt *a priori* hypotheses for these exploratory analyses, and interpretation was based on the estimated effects and their associated uncertainty rather than formal significance testing.

## METHODS

### Ethical approval and participants

The DXA datasets used for this study were drawn from four different resistance training interventions approved by the Auburn University Institutional Review Board. These studies are described herein as “YM, 10wk” (IRB approval # 20-136), “Younger, 10wk” (IRB approval # 19-249), “Older, 10wk” (IRB approval # 19-249), and “Older, 12wk” (IRB approval # 24-863). Each study was conducted in accordance with the latest version of the *Declaration of Helsinki*. Two trials (YM, 10wk and Older, 12wk) were not pre-registered as clinical trials. All participants provided verbal and written consent before beginning research procedures. Additionally, the Alabama Department of Public Heath (ADPH) approved all DXA scanning procedures for each study, and participants were screened to ensure that he/she did not exceed the acceptable annual radiation exposure limit established by ADPH for voluntary research studies. The number, sex, age, and body mass index of included participants are presented in Table 1.

**Table 1.** Pre-training phenotype data.

| Study | Participant # (sex) | Age (years) | BMI (kg/m<sup>2</sup>) |
| --- | --- | --- | --- |
| YM, 10wk | 31 (all M) | 23 ± 4 | 24.1 ± 3.0 |
| Younger, 10wk | 47 (13 M, 34 F) | 21 ± 2 | 23.1 ± 3.1 |
| Older, 10wk | 15 (6 M, 9 F) | 59 ± 5 | 31.8 ± 5.8 |
| Older, 12wk | 26 (12 M, 14 F) | 57 ± 7 | 28.1 ± 3.7 |
| All participants | 119 (62 M, 57 F) | 34 ± 18 | 26.0 ± 4.7 |

Legend: data show participant number (sex) as well as mean (± standard deviation) pre- intervention ages and body mass index values for included participants.

### Study designs for each intervention

Participants with minimal prior resistance training experience completed 10-12 weeks of whole-body resistance exercise performed twice per week for each of the four interventions utilized for this analysis. Exercises completed in each workout included hex bar deadlifts, 45º leg press, seated leg extensions, lying hamstring curls, machine chest press, and cable pulldowns. Training sessions were supervised by laboratory personnel with prior experience in instructing exercise techniques, and who ensured proper form and intensity were achieved. Training intensity was monitored using repetitions in reserve (RIR) as described by Zourdos et al. (16). RIR was recorded after every set of each exercise and load adjustments were made to achieve an RIR of 0-2. If RIR was >2, the load was increased by an additional ∼4.5-9 kg (10-20 pounds) for lower-body exercises and ∼2.3-4.5 kg (5-10 pounds) for upper-body exercises. If participants were unable to complete the prescribed number of repetitions, the load was decreased by ∼4.5-9 kg (10-20 pounds) for lower-body exercises and ∼2.3-4.5 kg (5-10 pounds) for upper-body exercises. All training programs utilized linear progression that increased load and decreased volume throughout the 10-12-week period. Moreover, training load was documented for every participant at each testing session to determine total volume-load over the course of the intervention.

The Younger, 10wk, Older, 10wk, and Older, 12wk studies involved supplementation paradigms. The two former studies involved approximately half the participants consuming a daily peanut protein supplement throughout the training intervention, whereas the other participants did not consume a supplement. In the Older, 12wk study, half of the participants consumed a reduced nitrate beetroot juice supplement daily during the intervention period, and half of the participants consumed the same volume of nitrate-rich beet root juice providing 13 mmol nitrates per day. We examined the pre-to-post intervention changes in total appendicular (dual-arm + dual-leg) bone-free lean tissue masses between supplementation paradigms for each study (Table 2) and confirmed that no differences existed between supplementation and control groups. More in-depth experimental procedures regarding each study can be found in prior publications for the YM, 10wk study (17), Older, 10wk study (18), and Younger, 10wk study (19). Note, while DXA outcomes are presented herein for the Older, 12wk study, beetroot supplementation effects on outcomes unrelated to the current research question (i.e., endothelial function and muscle tissue and fiber characteristics) are currently being drafted for publication.

**Table 2.** Supplementation effects on appendicular lean tissue mass changes.

| Study | $\Delta$ DXA app. bone-free lean mass (control group) | $\Delta$ DXA app. bone-free lean mass (supplementation group) | p-value |
| --- | --- | --- | --- |
| YM, 10wk | N/A (no supplementation) (n = 31) |  | --- |
| Younger, 10wk | 789 ± 907 g (n = 22) | 773 ± 717 g (n = 25) | 0.948 |
| Older, 10wk | 311 ± 1398 g (n = 8) | 697 ± 484 g (n = 7) | 0.453 |
| Older, 12wk | 542 ± 727 g (n = 13) | 584 ± 488 g (n = 13) | 0.862 |

Legend: data show that the nutritional supplementation interventions did not lead to statistically significant changes in DXA appendicular bone-free lean tissue mass changes when compared to placebo or no supplementation groups.

### Pre- and post-intervention anthropometrics and DXA

Participants arrived at the Auburn School of Kinesiology laboratory in the morning hours (7:00 to 10:00 AM) after an overnight fast in most cases. In less than 10% of cases, participants arrived between 10:00 AM to 1:00 PM following at least a 4-hour fast. Participants first submitted a urine sample (∼5 ml) to assess urine specific gravity levels using a handheld refractometer (ATAGO; Bellevue, WA, United States). Notably, all participants included in this analysis presented values less than 1.020 indicating that they were well-hydrated. Younger females also provided negative urine-based pregnancy tests to ensure that they were eligible for DXA scans according to IRB procedures. Participant height and body mass were then measured using a digital scale (Seca 769; Hanover, MD, USA). Next, participants underwent a whole-body dual DXA (Lunar Prodigy; GE Corporation, Fairfield, CT, USA). Associated software segmented with predefined regions of interest (ROI) were used to assess appendicular masses of bone as well as bone-free lean tissue masses. Additionally, a custom ROI tool enabled by the software was used to assess shoulder (acromion-to-acromion) and hip widths (greater trochanter-to-trochanter). Note that pre- and post-intervention testing occurred within a ±2-hour time differential to circumvent potential time of day testing variation. Likewise, post-intervention scans occurred ∼48-72 hours following the last resistance training bout. Though test-retest DXA scans were not performed on these participants, prior data from our laboratory test-calibrate-immediate retest on 10 participants produced an intra-class correlation coefficients of 0.998 for total body lean tissue mass [mean difference between tests (mean ± standard error) = 0.29 ± 0.13 kg] (20).

### Food log analysis

Participants were instructed to maintain their pre-intervention dietary habits throughout the interventions. For three of the four interventions, 4-day pre- and post-intervention food logs were administered to participants approximately one week prior to post-intervention testing whereby they were instructed to return the food logs to study staff. Participants were instructed to fill out logs in a detailed fashion for two typical weekdays and weekend days. Food logs were then inspected, and analysis was overseen by a registered dietician (A.D.F.). The Nutrition Data System for Research (NDSR; NDSR 2022; University of Minnesota) software was utilized by the study personnel to analyze Younger, 10wk and YM, 10wk food logs. The study personnel entered recalls into the Automated Self-Administered 24-hour Dietary Assessment (ASA24; National Cancer Institute, Bethesda, MD, USA) tool to analyze Older, 10wk study food logs. All nutritional data are expressed as average self-reported kilocalories (kcal) or grams of protein intake per day. Food logs were not collected in the Older, 12wk participants; however, again, these participants were instructed to maintain their pre-intervention dietary habits throughout the intervention.

### Volume-load calculation

Across all four studies, total volume-load was determined for each individual by calculating the sum of the per set volume-load (load used x repetitions performed) for all sets performed in the intervention.

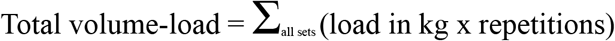

### Statistical analyses

All statistical analyses were performed using R version 4.4.0 (R Core Team, 2024). A Bayesian framework was adopted to enable flexible model specification, incorporate prior information, and provide transparent quantification of uncertainty. This approach allowed in-house estimates of measurement error for individual participants to be integrated directly into models, thereby improving the precision of parameter estimates. Posterior distributions allowed results to be interpreted in terms of probabilities, enhancing their practical interpretability.

To examine the relationship between baseline muscularity and hypertrophic responses, correlations between random intercepts (baseline values) and slopes (change scores) were estimated within hierarchical models. This approach accounted for regression to the mean—a common concern in pre-post designs—and enabled identification of individual-level associations. Given the limited number of time points (Pre and Post), informative priors were used to support model identifiability and to stabilize estimates. The potential influence of sex as a confounding variable was also explored, due to expected differences in baseline muscularity and responses to training.

Due to the strong correlations observed between pre-intervention appendicular bone characteristics and baseline lean tissue mass, conventional analyses of change scores were susceptible to issues of mathematical coupling and regression to the mean. To address this, associations between baseline skeletal features (e.g., appendicular bone characteristics, shoulder and hip widths) and appendicular lean tissue mass responses were examined by estimating correlations between participant-level random slopes (reflecting change over time) and baseline skeletal measures. This approach allowed baseline predictors to be related to the magnitude of adaptation without introducing spurious associations. Posterior samples of the correlation distribution were used to quantify the magnitude and uncertainty of these associations, providing a more robust and interpretable framework for assessing whether skeletal structure influenced training responsiveness. The same approach was used to evaluate associations between other participant attributes (e.g., age, training volume-load, self-reported energy intake) and lean tissue outcomes.

Finally, multivariable model development was guided by an iterative model comparison process using approximate leave-one-out cross-validation (LOO-CV). Starting from a well-specified base model, potential control variables including sex, age, body mass index (BMI), total training volume-load, and self-reported energy and protein intake were added individually and retained only if they improved model predictive performance as indicated by meaningful increases in expected log predictive density (ELPD-LOO). This approach helped to avoid overfitting while identifying variables with sufficient predictive contribution to warrant inclusion. Informed by this process, final models were used to explore whether baseline skeletal characteristics (e.g., shoulder and hip widths) moderated the appendicular lean tissue mass alterations with resistance training. Interaction effects were evaluated within the Bayesian framework, and interpretation focused on estimated effect sizes and their associated credible intervals rather than formal hypothesis testing.

## RESULTS

### Dual-arm bone characteristics and arm lean tissue mass training response

Figure 1 illustrates the relationship between baseline dual-arm bone-free lean tissue mass and the change in this variable with training. Posterior estimates suggested a weak positive association between baseline lean tissue mass and adaptation magnitude (*r* = 0.15 [75%CrI: –0.05 to 0.48, *p*(*r*>0) = 0.801]). However, this association attenuated after adjusting for sex (*r* = -0.15 [75%CrI: -0.40 to 0.18, *p*(*r*<0)=0.744]). Further analyses indicated that males exhibited a greater absolute response to training than females (Δ_Male:Female_ = -108 [75% CrI: -176 to -41 g, *p*(<0)=0.966]), as well as greater between-participant variability (σ_Male:Female_ = -139 [75%CrI: -193 to -87 g, *p*(<0)=0.998]).

**Figure 1.**
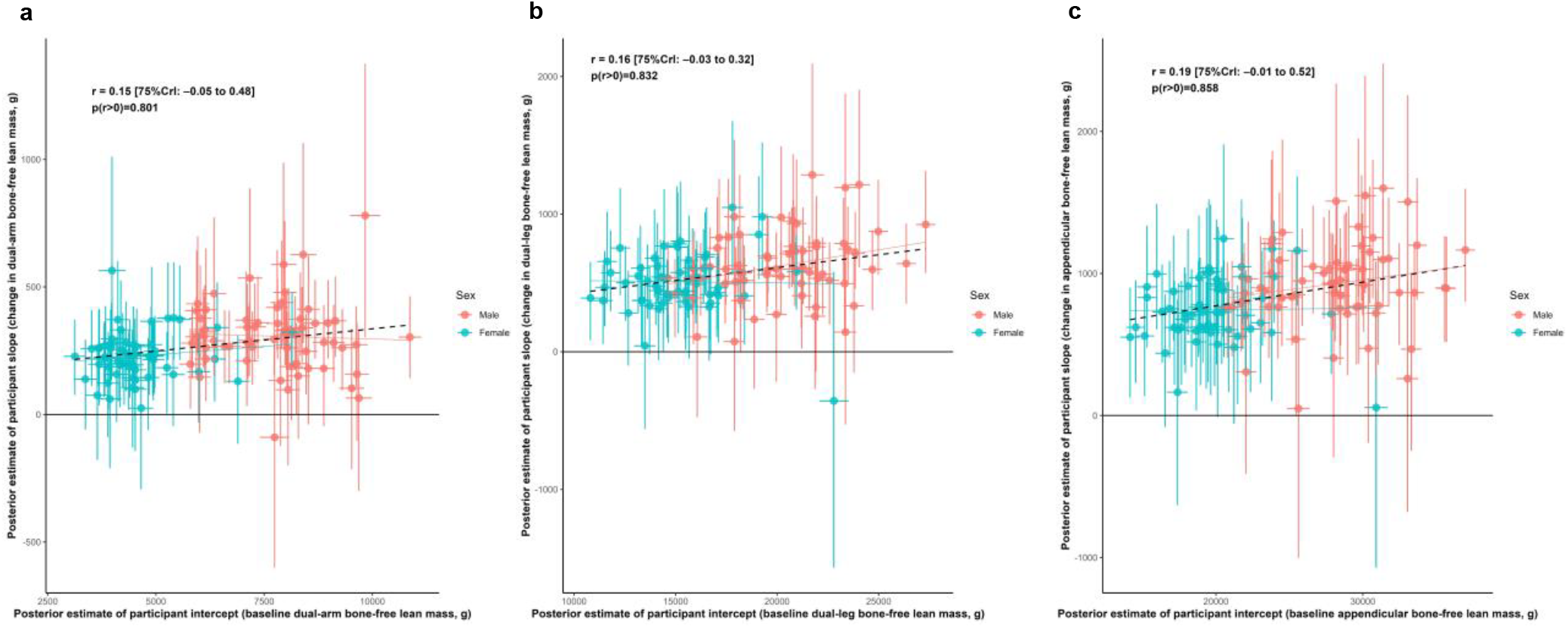
Associations between baseline lean tissue mass and training responses Legend: Main correlation estimates of the relationships between baseline and responses to training as represented by participant-specific intercepts and slopes, respectively. Panels include: A, dual-arm bone-free lean tissue mass; B, dual-leg bone-free mass; C, appendicular bone-free lean tissue mass. Error bars illustrate uncertainty in individual random effects as obtained from the posterior distributions. Solid lines represent sex-specific slopes, and dashed lines represented the slope for all participants. CrI: Credible interval.

### Dual-leg bone characteristics and leg lean tissue mass training response

Figure 1 also illustrates the relationship between baseline dual-leg bone-free lean tissue mass and the change in this variable with training. Posterior estimates suggested a weak positive association between baseline lean tissue mass and adaptation magnitude (*r* = 0.16 [75%CrI: – 0.03 to 0.32, *p*(*r*>0) = 0.832]). However, this association attenuated after adjusting for sex (*r* = 0.05 [75%CrI: -0.17 to 0.39, *p*(*r*<0) = 0.597]). Further analyses indicated that males exhibited a greater absolute response to training than females (Δ_Male:Female_ = -140 [75%CrI: -266 to -13 g, *p*(<0)=0.898]), as well as greater between-participant variability (σ_Male:Female_ = -68 [75%CrI: -168 to 31 g, *p*(<0) = 0.788]).

### Arm lean tissue mass, bone characteristics, and training response

Figure 2 illustrates relationships between baseline dual-arm bone characteristics and both baseline bone-free lean tissue mass and change in this variable to training. Posterior estimates indicated strong positive associations between baseline lean tissue mass and bone mineral content (*r* = 0.90 [75%CrI: 0.88 to 0.91, *p*(*r*>0) > 0.999]) as well as bone area (*r* = 0.64 [75%CrI: 0.58 to 0.69, *p*(*r*>0) > 0.999]). In contrast, the associations between hypertrophic response and these bone characteristics were weakly positive (BMC: *r* = 0.11 [75%CrI: -0.02 to 0.43, *p*(r>0) = 0.836]; bone area: *r* = 0.10 [75%CrI: -0.02 to 0.32, *p*(*r*>0) = 0.834]).

**Figure 2.**
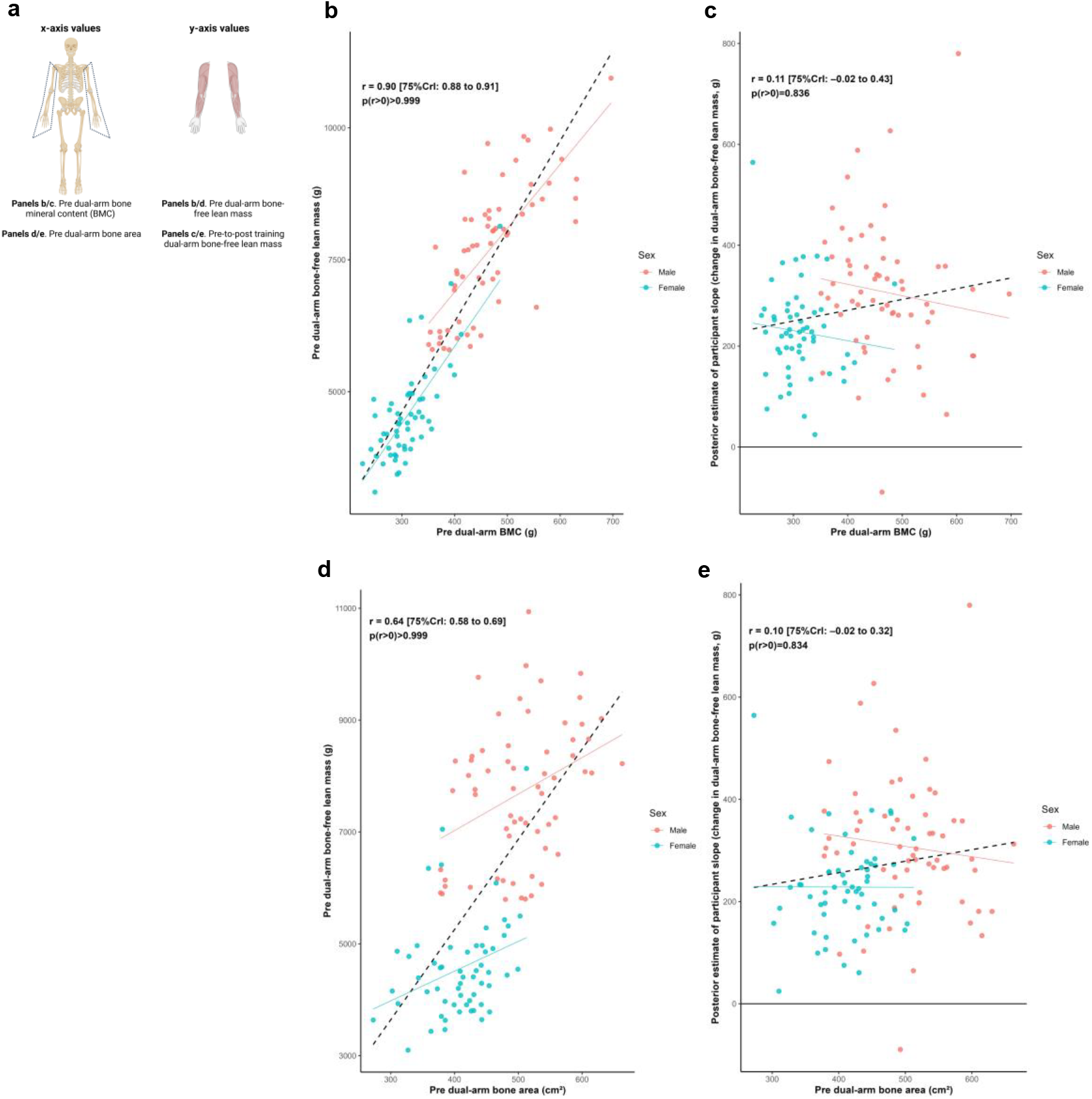
Dual-arm bone and bone-free lean tissue mass associations Legend: Panels include: A, schematic of the associations performed for this figure; B, pre-intervention dual-arm bone mineral content (BMC) versus pre-intervention dual-arm bone-free lean tissue mass; C, pre-intervention dual-arm BMC versus the training response in dual-arm bone-free lean tissue mass; D, pre-intervention dual-arm bone area versus pre-intervention dual-arm bone-free lean tissue mass; E, pre-intervention dual-arm bone area versus the training response in dual-arm bone-free lean tissue mass. Solid lines represent sex-specific slopes, and dashed lines represented the slope for all participants. CrI: Credible interval.

### Leg lean tissue mass, bone characteristics, and training response

Figure 3 illustrates relationships between baseline dual-leg bone characteristics and both baseline bone-free lean tissue mass and change in this variable to training. Posterior estimates indicated strong positive associations between baseline lean tissue mass and bone mineral content (*r* = 0.86 [75%CrI: 0.84 to 0.88, *p*(*r*>0) > 0.999]) as well as bone area (*r* = 0.84 [75%CrI: 0.82 to 0.87, *p*(*r*>0) > 0.999]). In contrast, the associations between hypertrophic response and these bone characteristics were weakly positive (BMC: *r* = 0.15 [75%CrI: 0.02 to 0.43, *p*(r>0) = 0.909]; bone area: *r* = 0.15 [75%CrI: 0.03 to 0.43, *p*(*r*>0) = 0.916]).

**Figure 3.**
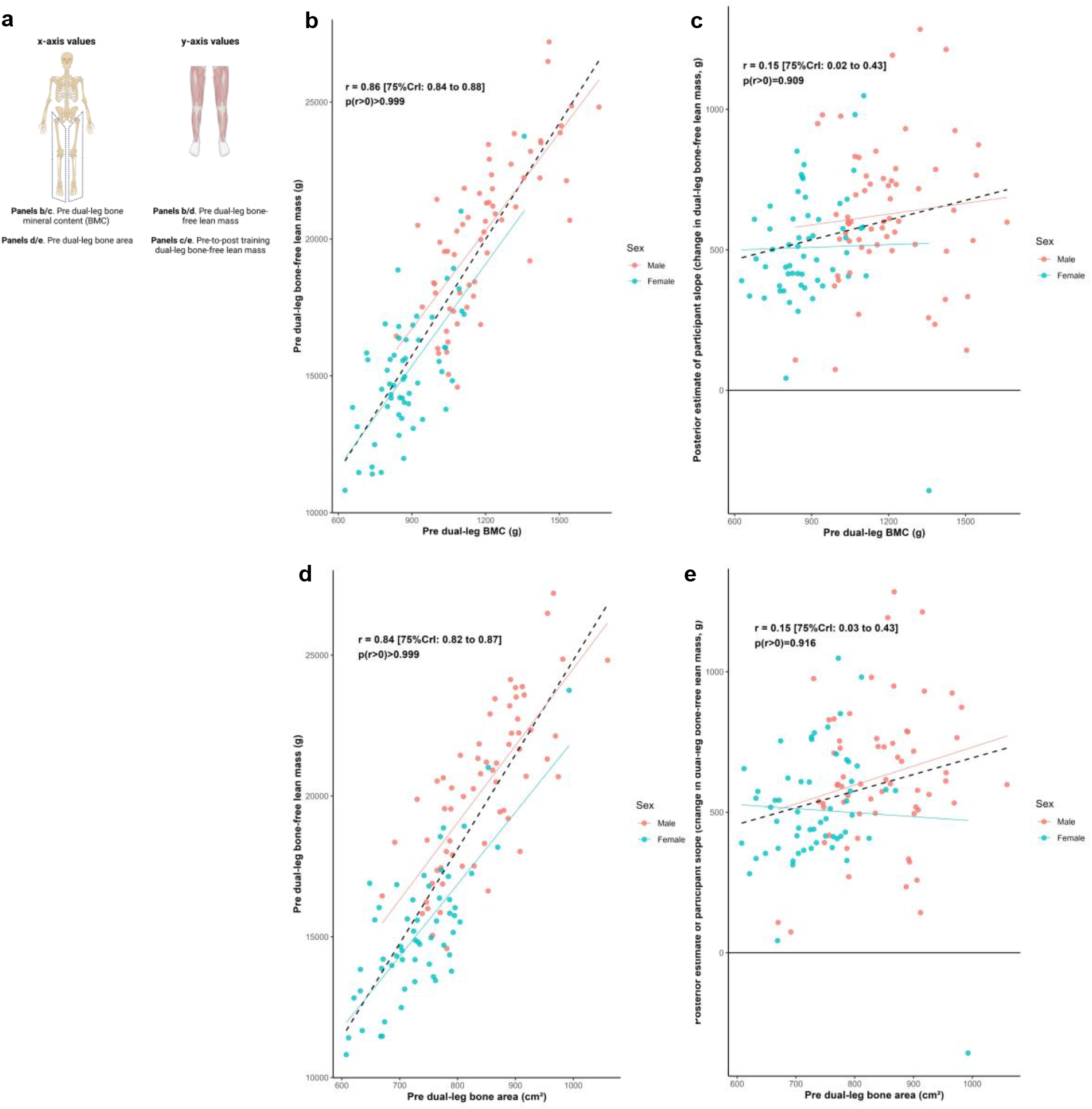
Dual-leg bone and bone-free lean tissue mass associations Legend: Panels include: A, schematic of the associations performed for this figure; B, pre-intervention dual-leg bone mineral content (BMC) versus pre-intervention dual-leg bone-free lean tissue mass; C, pre-intervention dual-leg BMC versus the training response in dual-leg bone-free lean tissue mass; D, pre-intervention dual-leg bone area versus pre-intervention dual-leg bone-free lean tissue mass; E, pre-intervention dual-leg bone area versus the training response in dual-leg bone-free lean tissue mass. Solid lines represent sex-specific slopes, and dashed lines represented the slope for all participants. CrI: Credible interval.

### Pelvic characteristics, leg lean tissue mass, and training response

Figure 4 illustrates the relationships between pelvic bone characteristics and both dual-leg baseline bone-free lean tissue mass and change in this variable to training. Posterior estimates indicated strong positive associations between baseline lean tissue mass and pelvic mineral content (*r* = 0.73 [75%CrI: 0.69 to 0.77, *p*(*r*>0) > 0.999]) as well as bone area (*r* = 0.74 [75%CrI: 0.70 to 0.78, *p*(*r*>0) > 0.999]). In contrast, associations between response to training and these bone characteristics were weakly positive (BMC: *r* = 0.11 [75%CrI: 0.00 to 0.37, *p*(r > 0) = 0.875]; bone area: *r* = 0.17 [75%CrI: 0.05 to 0.40, *p*(*r* > 0) = 0.929]).

**Figure 4.**
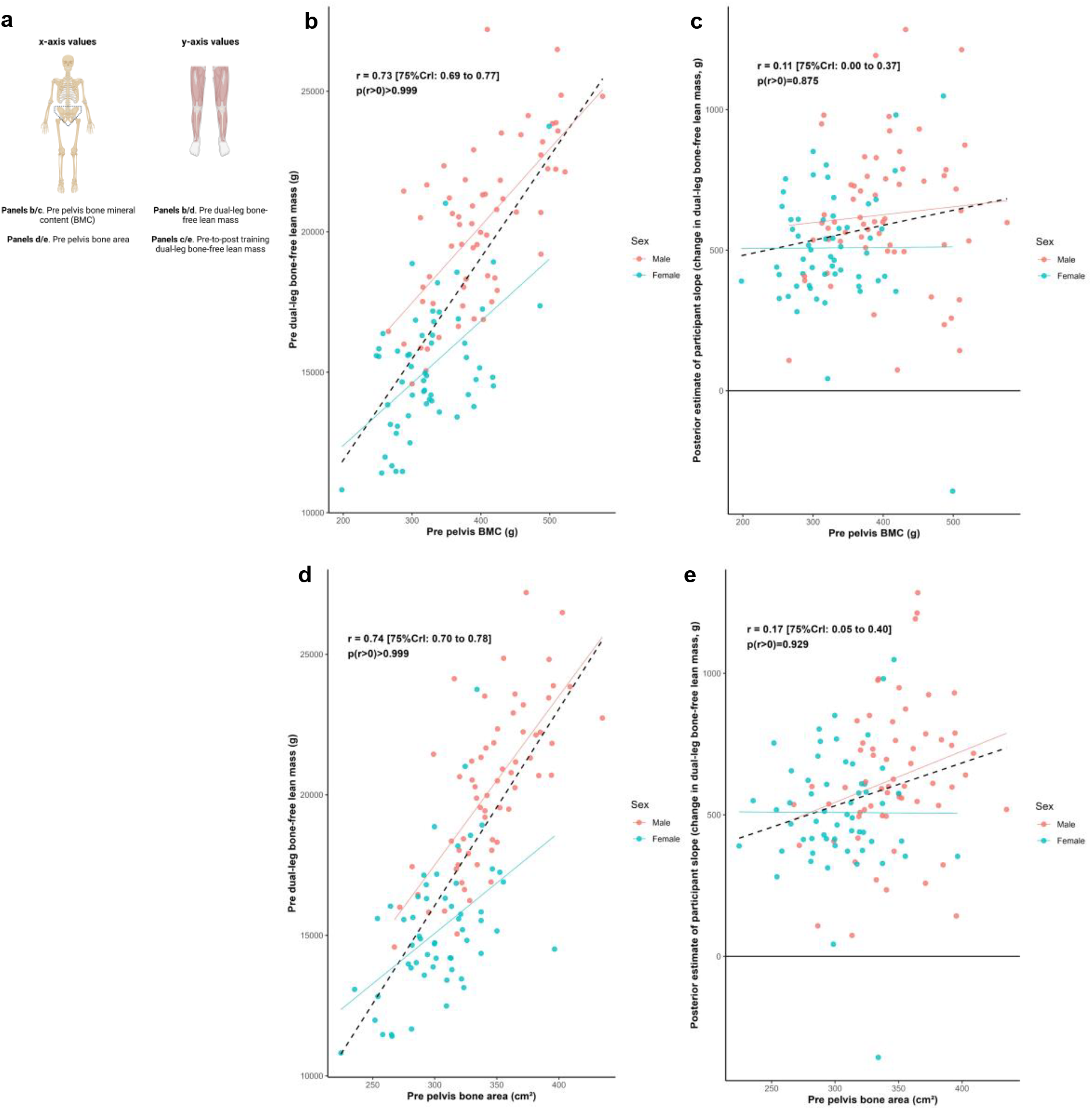
Pelvis and dual-leg bone-free lean tissue mass associations Legend: Panels include: A, schematic of the associations performed for this figure; B, pre-intervention pelvic bone mineral content (BMC) versus pre-intervention dual-leg bone-free lean tissue mass; C, pre-intervention pelvic BMC versus the training response in dual-leg bone-free lean tissue mass; D, pre-intervention pelvic bone area versus pre-intervention dual-leg bone-free lean tissue mass; E, pre-intervention pelvic bone area versus the training response in dual-leg bone-free lean tissue mass. Solid lines represent sex-specific slopes, and dashed lines represented the slope for all participants. CrI: Credible interval.

### Shoulder and hip widths, appendicular lean tissue mass, and training response

Figure 5 illustrates the relationships between shoulder and hip widths, and both dual-arm and dual-leg baseline bone-free lean tissue mass and response to training. Posterior estimates indicated a strong positive association between baseline dual-arm lean tissue mass and shoulder width (*r* = 0.76 [75%CrI: 0.72 to 0.79, *p*(*r*>0) > 0.999]), and a moderate positive association between baseline dual-leg lean tissue mass and hip width (*r* = 0.34 [75%CrI: 0.24 to 0.42, *p*(*r*>0) > 0.999]). Associations between the training responses and these skeletal characteristics were weakly positive for shoulder width (*r* = 0.19 [75%CrI: 0.06 to 0.41, *p*(r > 0) = 0.921]) and for hip width (*r* = 0.14 [75%CrI: 0.04 to 0.24, *p*(*r* > 0) = 0.925]). In contrast, there were limited differences in hip width across the sexes, but with slightly greater modeled improvements indicated with those with larger hip widths.

**Figure 5.**
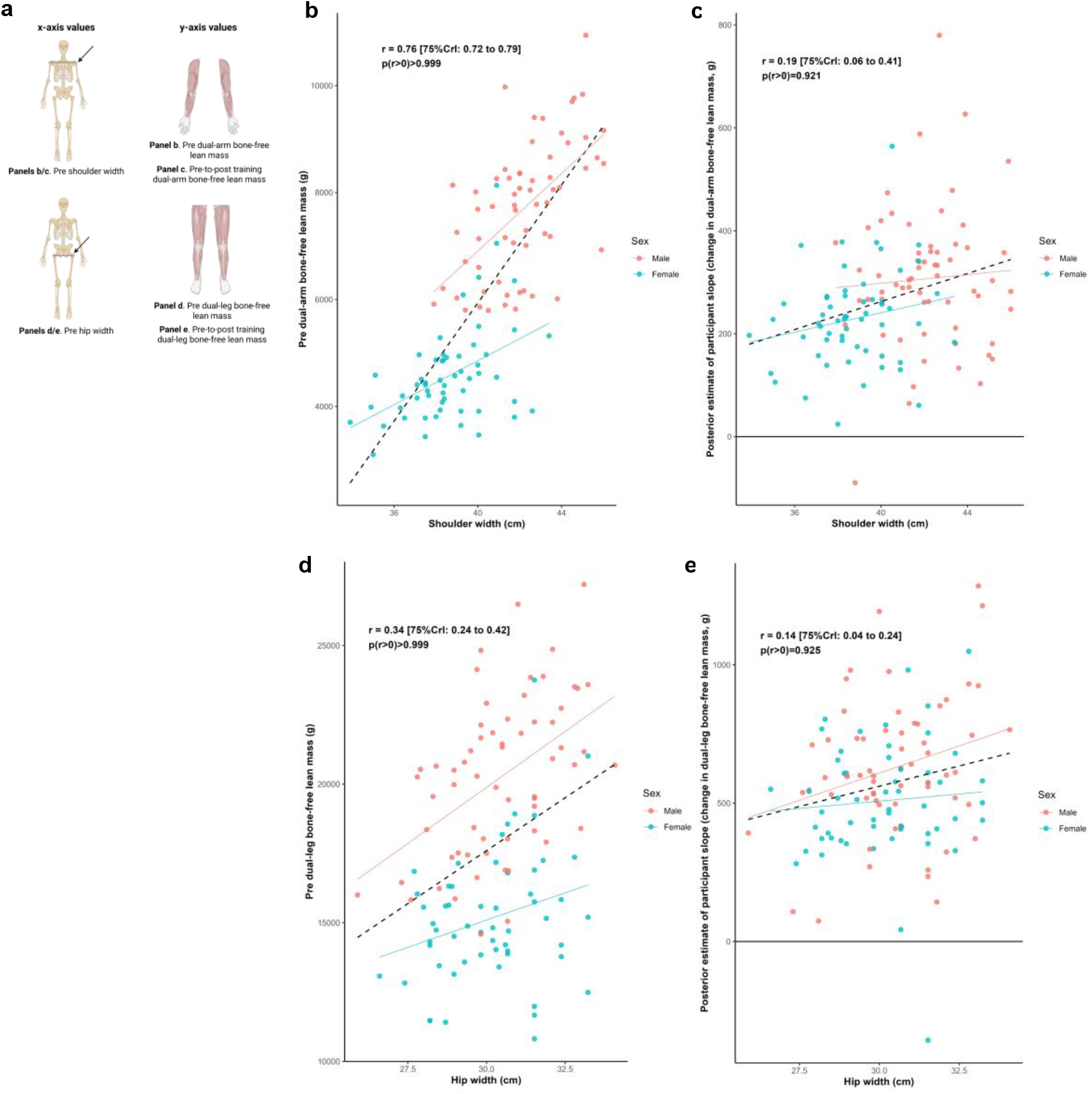
Skeletal width and appendicular bone-free lean tissue mass associations Legend: Panels include: A, schematic of the associations performed for this figure; B, shoulder width versus pre-intervention dual-arm bone-free lean tissue mass; C, shoulder width versus the training response in dual-arm bone-free lean tissue mass; D, hip width versus pre-intervention dual-leg bone-free lean tissue mass; E, hip width versus the training response in dual-leg bone-free lean tissue mass. Solid lines represent sex-specific slopes, and dashed lines represented the slope for all participants. CrI: Credible interval.

### Total appendicular lean tissue mass, other participant attributes, and training response

Figure 6 illustrates the relationship between the training response in total appendicular lean tissue mass and a range of participant attributes. Median estimates of correlations were positive (0.06≤ *r* ≤ 0.24) for training volume, self-reported energy intake, self-reported protein intake, and BMI; and negative for age (*r* = -0.08).

**Figure 6.**
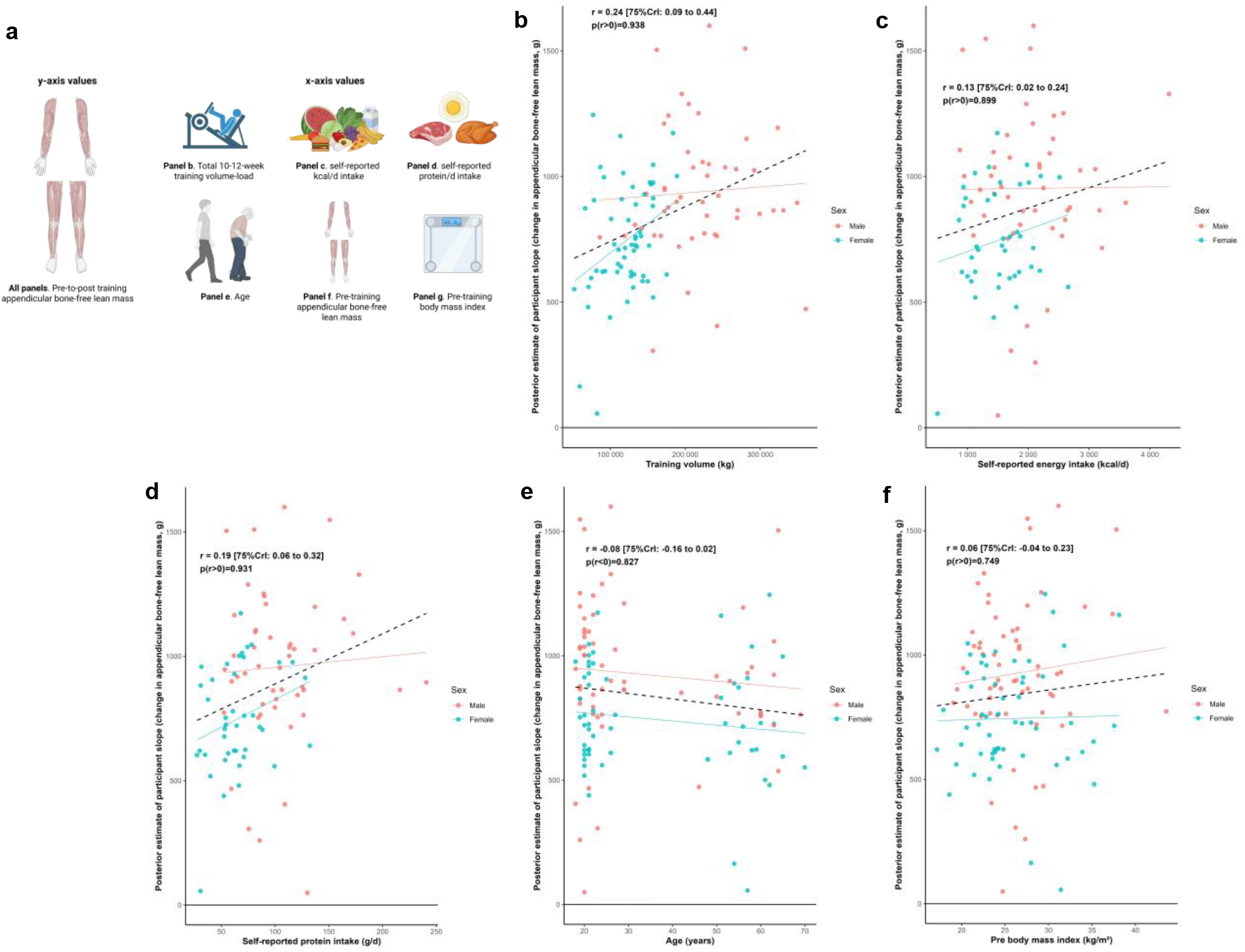
Other participant characteristic and appendicular bone-free lean tissue mass associations Legend: Panels include: A, schematic of the associations performed for this figure as well as the training response in total appendicular bone-free lean tissue mass versus volume-load during the resistance training intervention (B), self-reported energy intake (C) and protein intake (D) over a 4-day period during the last week of the intervention, age (E), pre-intervention total appendicular bone-free lean tissue mass (F), and pre-intervention body mass index (G). Solid lines represent sex-specific slopes, and dashed lines represented the slope for all participants. CrI: Credible interval.

Multivariable analyses examining whether shoulder and hip widths moderated changes in total appendicular lean tissue mass are presented in Table 3. In the initial model building phase, leave-one-out cross-validation indicated that none of the considered covariates except sex meaningfully improved predictive performance and were therefore excluded. After adjusting for sex, there was no evidence that either hip width (*β* = 0.9 [75%CrI: –5.5 to 7.1, *p*(*β* > 0) = 0.564]) or shoulder width (*β* = 1.0 [75% CrI: –3.7 to 5.8, *p*(*β* >0) = 0.597]) moderated changes in total appendicular lean tissue mass with training.

**Table 3.** Summary of model building and multivariable analysis exploring potential moderating effects of baseline hip and shoulder width on appendicular bone-free lean tissue mass changes.

| Variable | ELPD-LOO difference (standard error) | Regression interaction coefficient (75% CrI) | Bayesian posterior probability $p(>0)$ |
| --- | --- | --- | --- |
| **Univariable predictors** |  |  |  |
| Sex | 3.9 (1.8) |  |  |
| Training volume | -6.2 (3.0) |  |  |
| Self-reported kcal/d intake | -3.5 (0.8) |  |  |
| Self-reported protein g/d intake | -2.8 (1.1) |  |  |
| Age | 0.1 (0.8) |  |  |
| BMI | 1.3 (1.0) |  |  |
| **Multivariable predictors** |  |  |  |
| Sex + | -6.6 (2.8) |  |  |
| Shoulder width |  | 1.0 (-3.7 to 5.8) | 0.597 |
| Sex + | -4.5 (2.6) |  |  |
| Hip width |  | 0.9 (-5.5 to 7.1) | 0.564 |

Legend: Negative ELPD-LOO difference values indicate lower expected out-of-sample predictive performance compared to the base model. CrI: Credible interval.

## DISCUSSION

Our primary aim was to determine if pre-intervention bone characteristics were associated with resistance training-induced changes in lean tissue mass in a diverse cohort of participants. The resultant data suggest that, while pre-intervention bone measures were strongly associated with pre-intervention lean tissue mass outcomes, bone characteristics were not strongly associated with the appendicular lean tissue mass responses to 10-12 weeks of resistance training. Although weak positive correlations were observed in some cases (e.g., r ≈ 0.1 to 0.2), these appeared to be primarily driven by sex differences in baseline morphology and adaptation. A similar pattern was observed for baseline appendicular lean tissue mass, which showed a weak positive association with training-induced changes in this variable that was likewise attenuated after adjusting for sex. Our secondary analyses revealed that other attributes including total volume-load, dietary factors, age, and pre-intervention body composition metrics were only weakly associated with training-induced changes in appendicular lean tissue mass when individual correlations were performed. Finally, after adjusting for sex, multivariable analyses revealed minimal evidence that skeletal characteristics moderated changes in appendicular lean tissue mass in response to resistance training.

The dissociation between individual baseline bone parameters and changes in appendicular lean tissue mass with training was somewhat unexpected given the well-established relationship between bone and muscle tissue. A likely explanation for these findings is that, although bone and muscle development are coordinated during postnatal and pubescent growth, the adaptive muscular response to resistance training likely operates through distinct mechanistic pathways. In this regard, it has been established that mechanical overload in humans and rodents stimulates mechanotransduction-related mechanisms in muscle that in turn stimulate protein synthesis, satellite cell activation, and ribosome biogenesis (10). Several independent lines of evidence have also indicated that the satellite cell proliferation and ribosome biogenesis responses are predictive of hypertrophic responsiveness induced through resistance training (21-25). Additionally, differential skeletal muscle miRNA expression between high- and low-responders has been reported (28, 29). Changes in androgen receptor protein content have also been correlated with changes in fCSA (30, 31); however, our lab has reported that androgen receptor protein content decreases in both high and low responders to training (32). Notably, these processes may be relatively independent of bone characteristics, particularly over the relatively shorter 10-12-week interventions employed in the studies included in our analyses.

Our secondary analyses also revealed insightful outcomes. First, weak associations existed between lean tissue mass responses and several modulators including total intervention volume-load, self-reported dietary intakes, and age. Although these findings may seem unexpected, they do agree with prior literature. For instance, a recent meta-analysis supports that older individuals (∼55-65 years old) can experience similar relative increases in muscle mass versus younger cohorts in response to resistance training (26). Although these responses may be impaired with advanced aging (e.g. >80 years old) (27), none of our participants fell within this age group. The weak associations between the total appendicular lean tissue mass responses to training and self-reported energy/protein intakes as well as total intervention volume-load are also not surprising given that similar findings have been previously reported by our group and others in younger and older adults (23, 25, 28). Collectively, these results continue to highlight the complexity of resistance training adaptations.

## Methodological Considerations

There are several limitations to this study. First, DXA provides an estimate of bone-free lean tissue mass, which includes body fluids as well as skeletal muscle tissue. Evidence indicates that changes in DXA-derived lean tissue mass only moderately correlate with magnetic resonance-derived measurements of muscle cross-sectional area and/or volume (29, 30). Thus, caution should be employed when extrapolating lean tissue mass results to muscle hypertrophy. Second, although the 10-12-week intervention was sufficient to induce increases in lean tissue mass, it is possible that participants would have experienced additional increases had training been longer. Hence, the 10-12-week period may not have been long enough to completely parse our stronger associations between pre-intervention bone characteristics and hypertrophic outcomes. Third, our assessment of bone characteristics was limited to DXA-derived measures, which provide useful albeit incomplete bone data. Hence, future research examining more sophisticated imaging techniques (e.g., peripheral quantitative computed tomography or magnetic resonance imaging) could provide information regarding how bone geometry, cortical thickness, and/or trabecular architecture associate with hypertrophic potential. Finally, dietary data were assessed through self-reported food logs, and it is well known that these data can appreciably deviate from actual intakes (31).

### Conclusions

Individual pre-intervention bone characteristics, despite being strongly associated with pre-intervention lean tissue mass characteristics, are weakly associated with resistance training-induced measures of skeletal muscle hypertrophy. These findings continue to advance our understanding as to which physiological factors delineate hypertrophic responsiveness to resistance training.

## Acknowledgements

We thank the participants who volunteered and participated in the studies included in our analyses as well as other laboratory members who assisted with aspects of data collection unrelated to the current aims. Raw data related to the current study outcomes will be provided upon reasonable request by emailing the corresponding author.

